# Integrated phenotypic and metabolomic analyses identify elite chickpea haplotypes under water limitation

**DOI:** 10.64898/2026.07.15.738607

**Authors:** Nicholas Booth, Alexandra Cunningham, Crystal Sweetman, David A Day, Sunita A Ramesh

## Abstract

Chickpea (*Cicer arietinum*) is a globally important legume crop whose productivity is frequently constrained by terminal drought, particularly in rainfed systems. We used high-throughput phenotyping of 35 diverse chickpea haplotypes at contrasting watering regimes (80% and 40% field capacity) to identify superior haplotypes. Significant haplotype:watering interactions were observed for water-use dynamics, growth rates, biomass accumulation and nodulation, indicating strong genetic control over drought responses. Certain haplotypes (e.g., ICC2210 and ICC18839) maintained relatively high water-use efficiency and growth under stress, while others exhibited pronounced reductions in biomass and nodulation. Principal Component Analyses (PCA) were used to identify haplotypes associated with tolerant and sensitive stress phenotypes.

Metabolomic profiling revealed widespread reprogramming of metabolism under water limitation, with 57 of 82 metabolites significantly affected by treatment. A consistent decrease in tricarboxylic acid intermediates, including succinic acid, indicated altered energy metabolism, while accumulation of osmoprotectants such as proline and sucrose reflected adaptive responses to osmotic stress. Multivariate and ANOVA Simultaneous Component Analyses (ASCA) identified key metabolites as major contributors to haplotype-specific drought responses. These metabolites are linked to nitrogen metabolism, stress signalling and cellular protection mechanisms. These results demonstrate substantial variation in drought adaptation among chickpea haplotypes and confirm that the integration of phenotypic and metabolomic traits is a powerful approach to identify drought-resilient genotypes.

## INTRODUCTION

Chickpea (*Cicer arietinum*) is a legume grain crop and a rich source of protein, vitamins, minerals and fibre. Rising demand for plant-derived protein has driven increased global production of chickpea and other legumes. While India remains the largest producer of chickpea, Australia is the second largest producer and the biggest global exporter, with the majority of exports going to India, Bangladesh and Pakistan (Blauer 2023).

As with most crops, growth of chickpea is limited by a range of stresses, most notably drought, heat, acidity, salinity, Phytophthora root rot and Ascochyta blight. Water stress at different stages of growth in dry and arid zones leads to significant reduction in development and growth (Maqbool et al. 2017; Song et al. 2026). However, factors such as nutrient availability, soil pH, excessive nitrogen in the soil and extreme temperature may also affect nodulation, nitrogen fixation growth and yield (Zahran Hamdi 1999). The presence of endogenous soil nitrogen limits nodulation and nitrogen fixation in legumes generally (Streeter 1985), while an increase in temperatures above 30°C leads to delayed nodule initiation, development and nitrogen fixation in temperate legumes. Lower temperatures (<10 °C), on the other hand, reduce nitrogen fixation in tropical legumes (Ofosu-Budu et al. 1992; Lira Junior et al. 2005).

Chickpea establishes a symbiotic relationship with rhizobia to form nodules and fix atmospheric nitrogen (N) leading to decreased reliance on added fertiliser and improved growth. Expression of individual traits in haplotypes respond to changes in the environment (Bradshaw, 1965) and variation in nodulation and symbiotic effectiveness between cultivars has been observed in response to the water status of soils. The plasticity in traits such as seed number, seed size, number and size of plastids has been demonstrated across taxa (Mahto et al. 2025). In different legumes, nodule number and size at early developmental stages are affected by drought (Iqbal et al. 2022).

Major improvements to chickpea germplasm are needed to boost traits such as effective nodulation, nitrogen fixation and yield in response to changing climate, and sequencing of germplasm provides insights into the genetic variation. This genetic variation can be linked to phenotypic diversity for future breeding applications (Varshney et al. 2021).

Whole genome sequencing of 3,366 chickpea accessions (domesticated cultivars, landraces and wild types) by Varshney et al. (2021) identified several superior haplotypes for improvement related traits in chickpea landraces that could be incorporated into elite breeding lines following validation. We have compared thirty-five of these haplotypes imported from ICRISAT (India) that display superior yield traits using phenotypic screening and molecular analysis. The aim was to assess haplotype differences in nodulation, production of metabolites, response to oxidative stress and biomass when inoculated with the commercial strain of rhizobium, *Mesorhizobium ciceri* (CC1192).

## METHODS

### Plant growth and treatments

Thirty-five chickpea haplotypes were grown in a Smarthouse within the Australian Plant Phenomics Facility, University of Adelaide, Australia. Two watering regimes were established, based on field capacity (FC) of the sandy soil mix, control (80% FC) and stressed (40% FC). A split-unit design was employed to allocate five replicates of each haplotype and treatment (Supplementary Figure 1 for experimental design). Seeds were sown on 23 January 2025 (days after planting - DAP 0) in non-draining pots (200 mm diameter, 150 mm height) filled with sandy soil mix (sand: N40: clay loam in 2:5:3 ratio) with a pH of 7.6. Plants were grown under limiting nitrogen (0.5 mM KNO_3_) for 14 days before inoculating with 1 mL of *Mesorhizobium* strain CC1192, diluted in distilled H_2_O at an OD_550_ of 0.0236 (∼10^7^ rhizobia/plant) (Vincent 1970; Zaw et al. 2021). Plants were loaded on the conveyer system in the North East (NE) Smarthouse at the Plant Accelerator, Adelaide and automatically watered to weight every second day. Plants were imaged daily from 28 DAP and supplied with McNight’s Nutrient solution containing low nitrogen (0.5 mM KNO_3_) every 14 days until the end of the experiment. Water-stress was initiated 28 DAP by withholding water until 40% FC was achieved. Field capacity (100%) was calculated as the maximum weight of water held by the sandy soil mix under gravimetric conditions and equates to 16% soil moisture.

### Experimental design

The NE Smarthouse, fitted with conveyor systems and imaging stations (LemnaTec Scana-lyzer 3D) was used for the non-destructive, high-throughput phenotyping and occupied 21 Lanes (2-21) by 20 Positions (4–23). A split-unit design was employed to allocate five replicates of the 70 combinations of the haplotypes and watering regimes. Each Block was divided into 35 Main Units, with each Main Unit consisting of two adjacent Carts within a Lane; the 35 Main Units are arranged in seven rows (Lanes) by 5 columns (5 pairs of Main Units in the same pair of Positions). The main-unit design, used to allocate the Lines, was a near-A optimal, resolved, latinized, row-column design comprised of five Blocks, each occupying 7 Lanes by 10 Positions. The two waterings were randomized to the two Carts within each Main Unit. The main-unit design was obtained using the R package *odw* (Butler, 2022), a package for the R statistical computing environment (R Core Team, 2025). Then, the full design was randomized using the R package *dae* (Brien, 2024). The randomized layout of the experiment on the conveyor system in the NE Smarthouse is supplied in Supplementary Figure 1.

### Plant growth analysis

Shoot and water use traits were obtained from plant imaging data on the Smarthouse conveyor system, and harvest data obtained after imaging. The raw imaging data was processed using the Smoothing and Extraction of Traits (SET) method described by Brien et al. (2020) to produce further shoot and water use traits, as well as twenty-eight per-plant responses or features. Calculations were carried out using growthPheno (Brien, 2025c), a package for the R statistical computing environment (R Core Team, 2025).

### Shoot traits

Projected Shoot Area (PSA, kpixels) was calculated for each plant and imaging DAP as the sum of the number of plant pixels in the images from three camera views, comprising two side views and a view from above. The relative PSA growth rate (PSA RGR Day^-1^) was calculated for each plant and imaging DAP, except DAP 27, by differencing consecutive PSA and ln(PSA) values, respectively, and dividing by their DAP differences. Following exploratory smoothing of the PSA using the traitSmooth function from growthPheno, the trait smoothed PSA (sPSA kpixels) was obtained for each plant and imaging DAP using segmented, logarithmic smoothing of the PSA that involved separate smoothing of the two segments corresponding to the pre-and post-leaf-harvest periods, namely DAPs 27-49 and DAPs 50-62. For each period, smoothing employed P-splines with the smoothing penalty set to ten. The smoothed growth-rate traits, sPSA AGR (kpixels) and sPSA RGR (day^-1^), were computed from the sPSA analogously to the PSA AGR and PSA RGR.

### Water usage traits

Pot weights before and after watering were recorded for each watering DAP. Water use (WU mL) was defined as the net change in pot weight between successive watering days and adjusted for applications of McNight’s Nutrient solution on DAP 28 and 50. This was converted to daily water use rate (WUR mL day^-1^) by dividing by the DAP differences. Exploratory smoothing of the WUR using traitSmooth resulted in the production of the trait smoothed WUR (sWUR mL day^-1^) that was obtained by direct smoothing the WUR using P-splines with the smoothing penalty set to three. The trait smoothed Water Use Index (sWUI kpixels mL^-1^) is defined as the ratio of the sPSA AGR to the sWUR. Higher values of sWUI correspond to higher water efficiency. No attempt was made to differentiate between biological consumption and evaporative loss.

### Harvest traits

Two leaves were harvested from each plant after imaging on 49 DAP and snap frozen in liquid N_2_. Whole plants were harvested 63 DAP and dissected into shoots, roots, and nodules. Shoot, root and nodule fresh weights (FW, g) were recorded on the day of harvest, while shoot and root dry weights (DW, g) were measured after drying at 70°C for 48 h.

### Malondialdehyde (MDA)

MDA equivalents were measured using the Thiobarbituric Acid Reactive Substances (TBARS) assay, as described by Singh et al. (2014). Homogenised frozen leaf tissue (50 mg) was extracted in 5% (w/v) trichloroacetic acid (TCA), combined in equal volumes with 20% (w/v) TCA +/-0.01% (w/v) butylated hydroxytoluene and heated to 96°C for 30 minutes. The absorbance of the reaction mixture was measured at: 532 nm (TBA-MDA complex); 600 nm (nonspecific turbidity); and 440 nm (interfering soluble sugars) (Hodges et al. 1999). MDA equivalents were calculated using the modified equation proposed by Du and Bramlage (1992) and compared against a standard curve.

### GC-MS

Untargeted metabolomics using Gas Chromatography-Mass Spectroscopy (GC-MS) was performed at Flinders Analytical, Flinders University using a method modified from (Howell et al. 2009; Shingaki-Wells et al. 2011). Metabolites were extracted from 30 mg of freeze-dried leaf tissue in 300 µL of ice-cold metabolite extraction medium consisting of 85% (v/v) HPLC-grade methanol, 15% (v/v) deionized water, and ribitol (5 ng µL^-^¹) as an internal standard. Samples were shaken at 700 rpm for 15 min at 70°C, centrifuged and the supernatant was reextracted in an equal volume of deionized water and two volumes of chloroform. The polar phase was dried under vacuum and derivatised in 20 µL methoxyamine hydrochloride (20 mg mL^-^¹ in anhydrous pyridine) at 30°C for 90 minutes prior to the incubation with 30 µL of N-methyl-N-(trimethylsilyl) trifluoroacetamide (MSTFA) at 37°C for 90 minutes. Derivatised samples were transferred to amber GC–MS vials fitted with low-volume inserts, and an n-alkane retention index calibration mixture was added prior to analysis. Samples were analysed using GC-MS Agilent 7890A/5975C fitted with a EZGuard J&W VF-5ms GC Column, 30 m, 0.25 mm, 0.25 µm, 10 m EZ-Guard and 7-inch cage (CP9013). GC-MS run conditions were as specified by Howell et al. 2009. Raw GC-MS data was processed using MSDIAL-v5 with peak identification against the KovatsRI-VS3 DB Public library (Tsugawa et al. 2015).

## Statistical analysis

To produce phenotypic estimated marginal means (EMMs) (Searle et al. 1980), each response was analyzed using the R packages ASReml-R (Butler et al. 2017) and asremlPlus (Brien and Brien 2020). The following maximal linear model was fitted to each response:

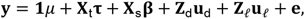

where **y** is the response vector of values for the response being analyzed; *μ* is the overall mean in this experiment for the response; **τ** is the vector for the fixed terms core to the experiment; **β** is the vector for fixed large-scale spatial terms; **u**_d_ is the vector for the random large-scale spatial terms arising from the design used; **u***_P_* is the vector for the random terms used in modelling the local spatial variation; **e** is the vector for the residual term; the vector 1 is the vector of ones; the matrices **X***_t_*, **X**_s_, and **Z**_d_ are the design matrices for the corresponding terms; and **Z***_l_* is the design matrix for the tensor-product splines or for an autocorrelation model of order one used in modelling the local spatial variation between individual plants.

For each response, Wald *F*-tests with *α* = 0.05 were used to test the haplotype and watering effects. Testing began with the interaction effect of haplotype and watering; if this was not statistically significant, tests were then performed for the haplotype and watering main effects. EMMs were produced that are based on the model term comprised of the core factors in the experiment, Line and Watering. Least significant differences for *α* = 0.05 [LSD (5%)] were calculated, from which error bars that are Least Significant Intervals (LSIs) were computed as an EMM ± half-LSD (5%).

Processed GC-MS data were analyzed using MetaboAnalyst 6.0 (Pang et al. 2024) or carried out directly using R packages tidyverse (Wickham et al. 2019) and factoextra (Kassambara and Mundt 2016). Biochemical changes in oxidative stress markers and metabolites were assessed in GraphPad Prism 10 and plotted as min to max box and whisker plot with statistical significance calculated through a two-way ANOVA with Šidák multiple comparisons test.

## RESULTS

### Haplotypes differ in water usage

Variability in the smoothed water use rate (sWUR) was evident amongst the 35 haplotypes at the time of drought initiation (Figure 1A). During the initial withholding of water (27-34 DAP), the sWUR of water-stressed plants varied little but trended towards a decrease. In contrast, Well-watered plants continued to increase in sWUR (Figure 1A). Clear separation in the sWUR of well-watered and water-stressed plants was evident from 34-44 DAP, where a steep decrease in sWUR of watered-stressed plants was observed until 57 DAP before increasing once more (Figure 1A). The sWUR of well-watered plants mostly increased with DAP (Figure 1A). The haplotype interaction on smoothed water use index (sWUI) was significant for all tested time intervals: at 34-44 DAP the watering interaction became significant, but the haplotype:watering interaction was non-significant (Figure 1B). For the remaining tested intervals, the haplotype:watering interaction was significant.

**Figure 1:**
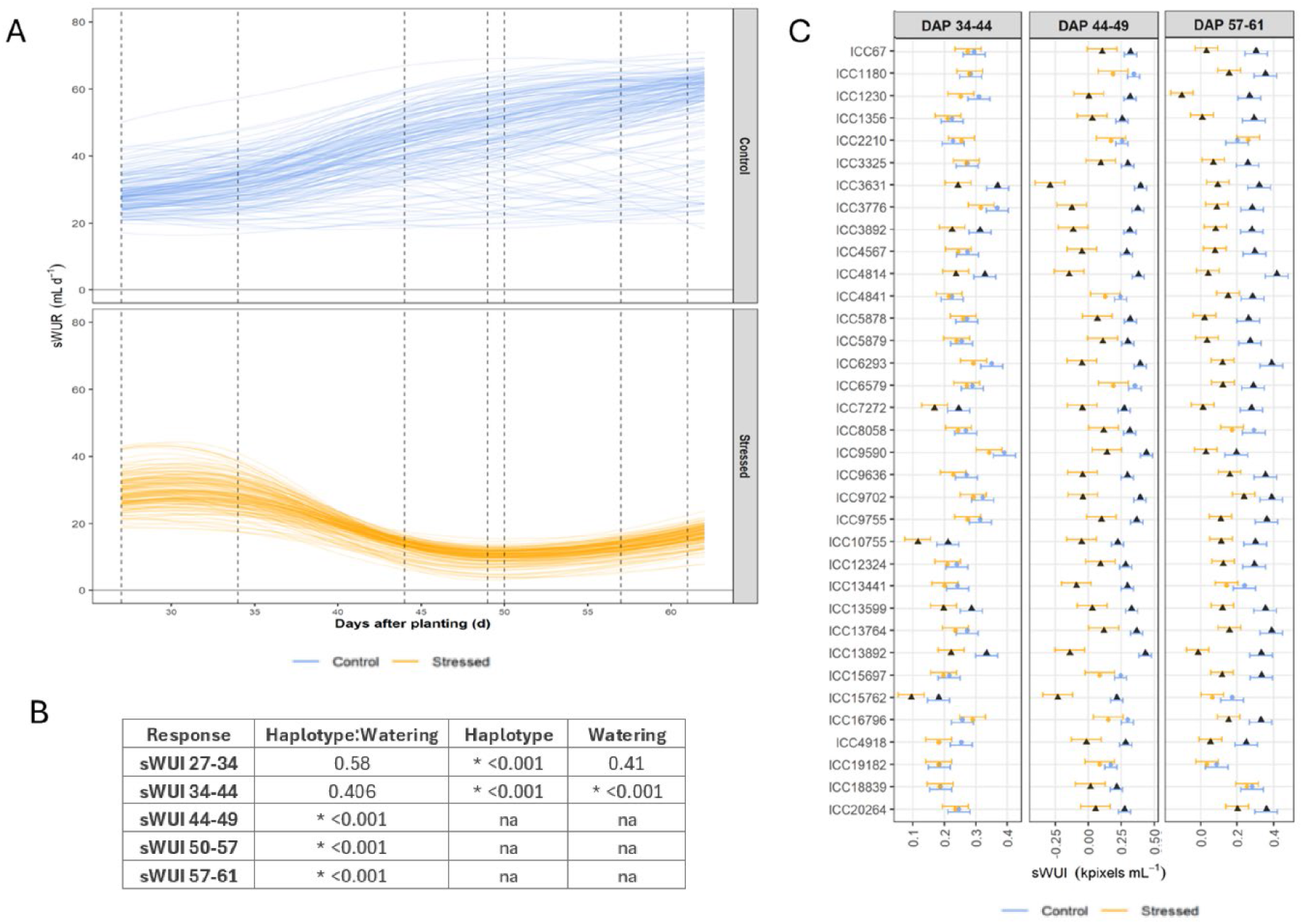
Water usage of the 35 *C. arietinum* haplotypes subjected to two water regimes. (A) Smoothed Water Use Rate (sWUR) from 27 to 62 Days after Planting (DAPs). Each curve corresponds to one plant. Vertical dashed lines correspond to the intervals used in the analysis. (B) The p-values of the Wald F-statistics that test the three core effects, the interaction of haplotype and watering and the haplotype and watering main effects. Statistically significant terms (p≤0.05) are flagged with an asterisk (*). An entry of na (not applicable) indicates that the effect was not tested because the haplotype:watering interaction is significant (p≤0.05). (C) Estimated marginal means (EMMs) for the smoothed Water Use Index (sWUI) for intervals covering Days after Planting (DAPs) 34-44, 44-49 and 57-61. Error bars are an EMM (n=5) ± half least significant difference (5%). The EMMs from the same DAP interval and watering level are significantly different (p≤0.05) if their error bars do not overlap. Haplotypes for which the difference between the watering treatments is significant (p≤0.05) are plotted with a solid triangle for both treatments.

Haplotype dependant sWUI was observed amongst the well-watered haplotypes: ICC15762 and ICC19182 had the lowest sWUI (<0.21 kpixel/mL), while high sWUI was more variable (Figure 1C). ICC9590 initially displayed high sWUI at 24-44 and 44-49 DAP (0.391 and 0.442 kpixel/ml) but decreased at 57-61 DAP (0.195 kpixel/mL). ICC4814, ICC6293 and ICC9702 haplotypes generally displayed a high sWUI during development (>0.33). Responses to the watering treatments at 34-44 DAP were significant for seven of 35 haplotypes, increasing drastically under prolonged water limitation to 28 significant interactions at 44-49 DAP (Figure 1C). Prior to sampling, at 57-61 DAP, the sWUI of 29 haplotypes was significantly different between well-watered and water-stressed plants. The haplotypes not significantly impacted by water limitation include: ICC2210; ICC8058; ICC13441; ICC15762; ICC19182; and ICC18839 (Figure 1C). ICC2210 and ICC18839 maintained a high sWUI under both well-watered and water-stressed conditions, ranging from 0.188-0.282 kpixel/mL for well-watered and 0.173-0.262 kpixel/mL for water-stressed conditions, respectively, across the tested intervals (Figure 1C). ICC8058 and ICC13441 both had a high sWUI under well-watered conditions, but this was highly variable under water-stressed conditions. ICC15762 and ICC19182 both displayed low sWUI across the tested intervals under well-watered (0.173-0.200 and 0.0869-0.184 kpixel/mL, respectively) and water-stressed conditions (-0.233-0.0948 and 0.0347-0.183 kpixel/mL, respectively) (Figure 1C).

### Haplotype growth over time under water limitation

At 27-34 DAP, the smoothed projected shoot area (sPSA) and relative growth rate (RGR) between well-watered and water-limited plants were not significantly different between any of the 35 haplotypes (Figure 2). A statistically significant interaction was observed between haplotypes for this time interval (Figure 2B). At 34-44 DAP, the sPSA RGR decreased for all plants with both haplotype and watering interactions becoming significant (Figure 2). The effect of watering regime on the sPSA RGR was non-significant for only haplotypes ICC2210 and ICC16796. At 44-49 and 50-57 DAP, the sPSA RGR slowed, with haplotype:watering interactions becoming significant (Figure 2). The data at 49-50 DAP included the removal of a leaf sample on 49 DAP that led to a discontinuity in sPSA AGR, and was therefore excluded from the analysis.

**Figure 2:**
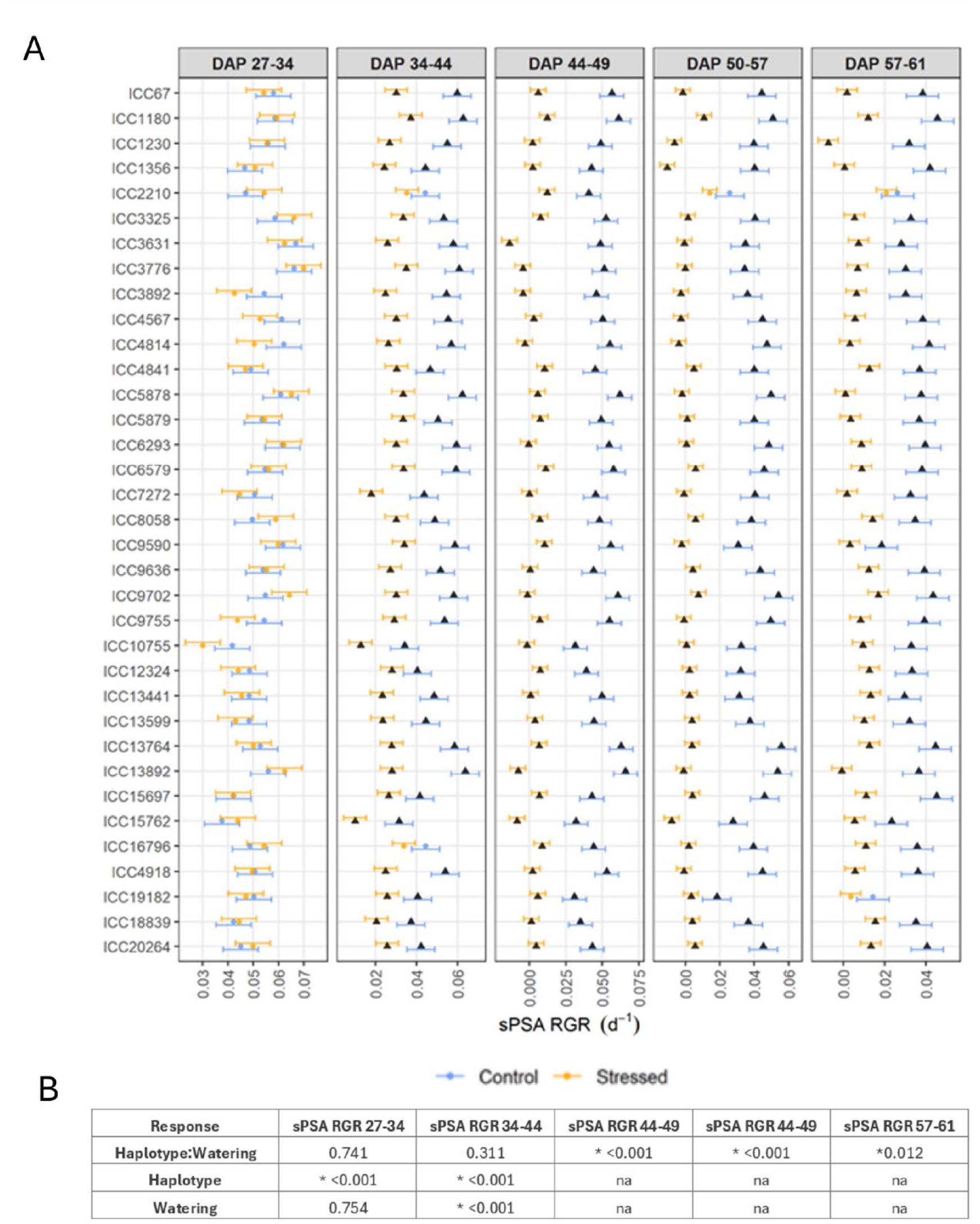
Estimated marginal means (EMMs) for the Relative Growth Rates of the smoothed Projected Shoot Area (sPSA RGR) for 35 *C. arietinum* haplotypes subjected to two water regimes. Intervals covering Days after Planting (DAPs) 27-34, 34-44, 44-49, 50-57 and 57-61 correspond to the intervals used in the analysis (n=5). (A) Error bars are an EMM ± half-least significant difference (5%) for the Relative Growth Rates of the smoothed Projected Shoot Area (sPSA RGR). The EMMs from the same DAP interval and Watering level are significantly different (p≤0.05) if their error bars do not overlap. Lines for which the difference between the Watering treatments is significant (p≤0.05) are plotted with a solid triangle for both treatments. (B) The p-values of the Wald F-statistics that test the three core effects, the interaction of haplotype and watering and the haplotype and watering main effects. Statistically significant terms (p≤0.05) are flagged with an asterisk (*). An entry of na (not applicable) indicates that the effect was not tested because the haplotype:watering interaction is significant (p≤0.05).

At 57-61 DAP, the well-watered plants continued to exhibit a decrease in the sPSA RGR while water-limited plants remained largely constant but the haplotype:watering interaction was significant (Figure 2). The final growth interval had two non-significant watering interactions in ICC2210 and ICC1912. In both cases, the sPSA RGR of well-watered plants was low (0.0263 and 0.0143 d^-1^, respectively). Four other haplotypes also displayed low sPSA RGR (<0.03 d^-1^) under well-watered conditions: ICC3631, ICC9590, ICC13441 and ICC15762. Seven haplotypes maintained high sPSA RGR (>0.04 d^-1^) under well-watered conditions over the final growth period: ICC1180, ICC1356, ICC4814, ICC9702, ICC13764, ICC15697 and ICC20264 (Figure 2A). Under water-limited conditions the sPSA RGR was much lower, with only three haplotypes (ICC2210, ICC9702 and ICC18839) maintaining a growth rate >0.015 d^-1^, whereas four haplotypes (ICC1230, ICC13892, ICC1356 and ICC5878) had growth rates <0.001 d^-1^ (Figure 2). This is reflected in the sPSA (kpixels) measurements of Supplementary Figure 2.

### Variation in total plant biomass at harvest

Following 28-days of treatment, well-watered and water-limited plants were harvested for shoot, root and nodule biomass measurements (Figure 3). The haplotype:watering interaction was significant for both shoot DW and nodule FW but not significant for root DW (Figure 3). Individual interactions between both haplotype and watering regime were significant for root DW.

**Figure 3:**
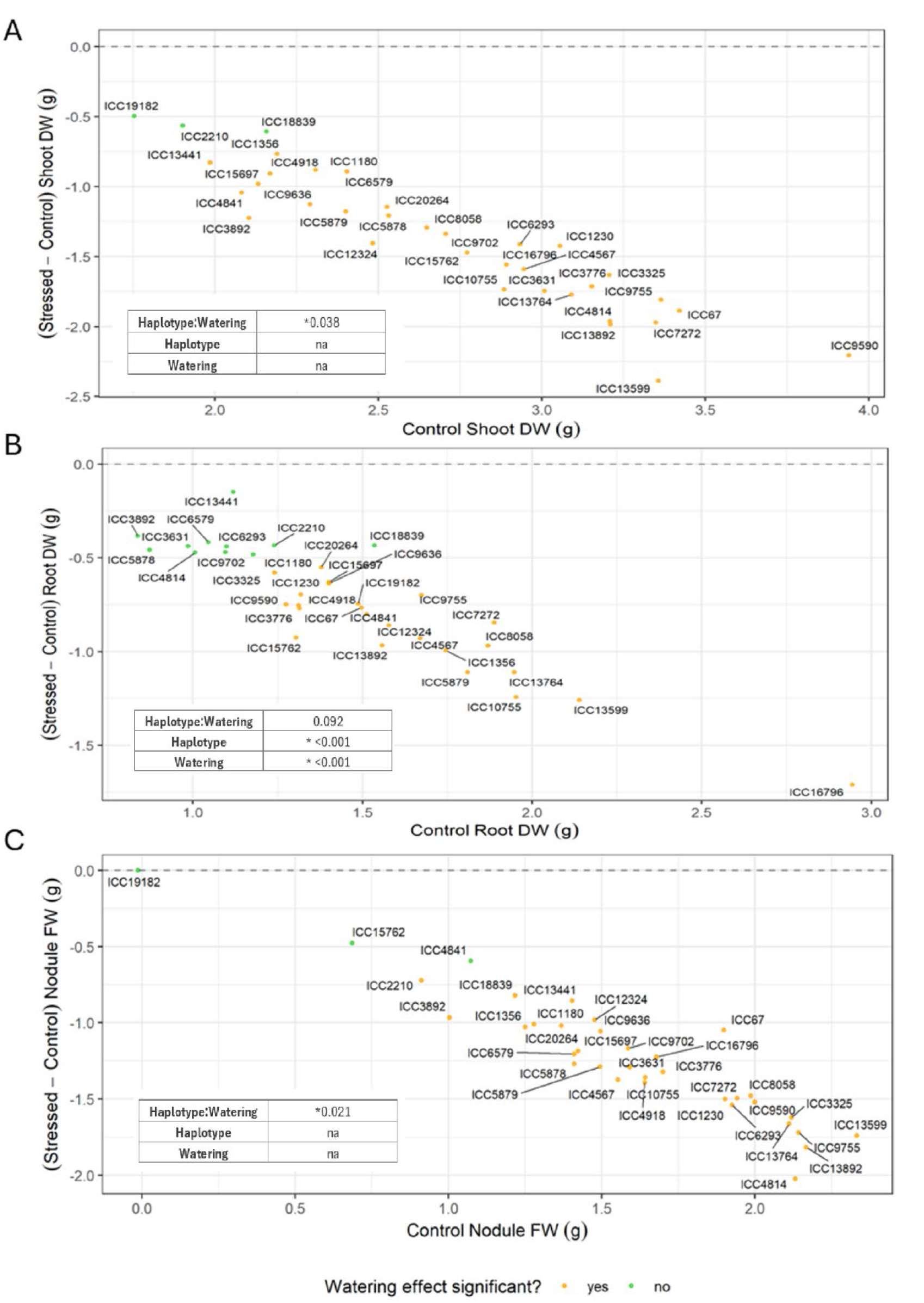
Scatterplot of the Stressed minus Control estimated marginal means (EMMs) vs Control EMMs for 35 *C. arietinum* haplotypes for (A) Shoot Dry Weight, (B) Root Dry Weight and (C) Nodule Fresh weight at the termination of experiment. Orange indicates a significant difference (p≤ 0.05) between the Stressed and Control treatments; green indicates the difference is not significant (n=5). The p-values of the Wald F-statistics that test the three core effects, the interaction of Haplotype and Watering and the Haplotype and Watering main effects. Statistically significant terms (p≤0.05) are flagged with an asterisk (*). An entry of na (not applicable) indicates that the effect was not tested because the haplotype:watering interaction is significant (p≤0.05).

Haplotype dependency was observed across shoot DW, ranging from 3.94 to 1.75 g per plant under well-watered, and 0.879 and 1.73 g per plant under water-limited conditions (Figure 3A; Supplementary Figure 3A). Three haplotypes did not display a significant difference in shoot biomass after 28-days of treatment: ICC2210, ICC19182, ICC18839 (Figure 3A).

Several haplotypes (11 of 35) showed no significant difference in root biomass following treatment (Figure 3B). In ICC2210 and ICC18839 both shoot and root biomass were not impacted (Figure 3A/B).

Drought had no significant impact on nodulation in ICC4841 and ICC15762, but both showed poor nodulation compared to the nodule FW of other screened haplotypes. ICC19182 is a non-nodulating haplotype and therefore displayed 0.00 g of nodule FW. ICC13599 and ICC13892 had the highest nodulation under well-watered conditions (>2.15 g plant^-1^). Furthermore, ICC13599 sustained a high level of nodulation (>0.590 g plant^-1^) under water-stressed conditions, while ICC67 had the highest level of nodulation under drought (0.847 g plant^-1^) (Supplementary Figure 3C).

### Stress induced responses to water limitation

Phenotypic traits and oxidative stress measurements for the 35 haplotypes was summarised through principal component analysis (PCA) biplot (Figure 4). PC1 explained 44.7% of the total variation, while PC2 explained 23.6%, jointly accounting for 68.3% of the haplotype variation in water-limiting conditions relative to well-watered conditions. Traits associated with increased tolerance to water-limitation pushed haplotypes strongly negative on PC1, while a similar pattern was noted for PC2 but less pronounced (Figure 4). The resulting drought sensitivity index classified 20% of haplotypes mildly, 54% moderately and 26% severely impacted by water limitation (Table 1). These were broken down further by seed type, with 27%, 64% and 9% of Kabuli type haplotypes, and 20%, 54% and 26% of Desi haplotypes mildly, moderately and severely impacted by water limitation, respectively (Table 1).

**Figure 4:**
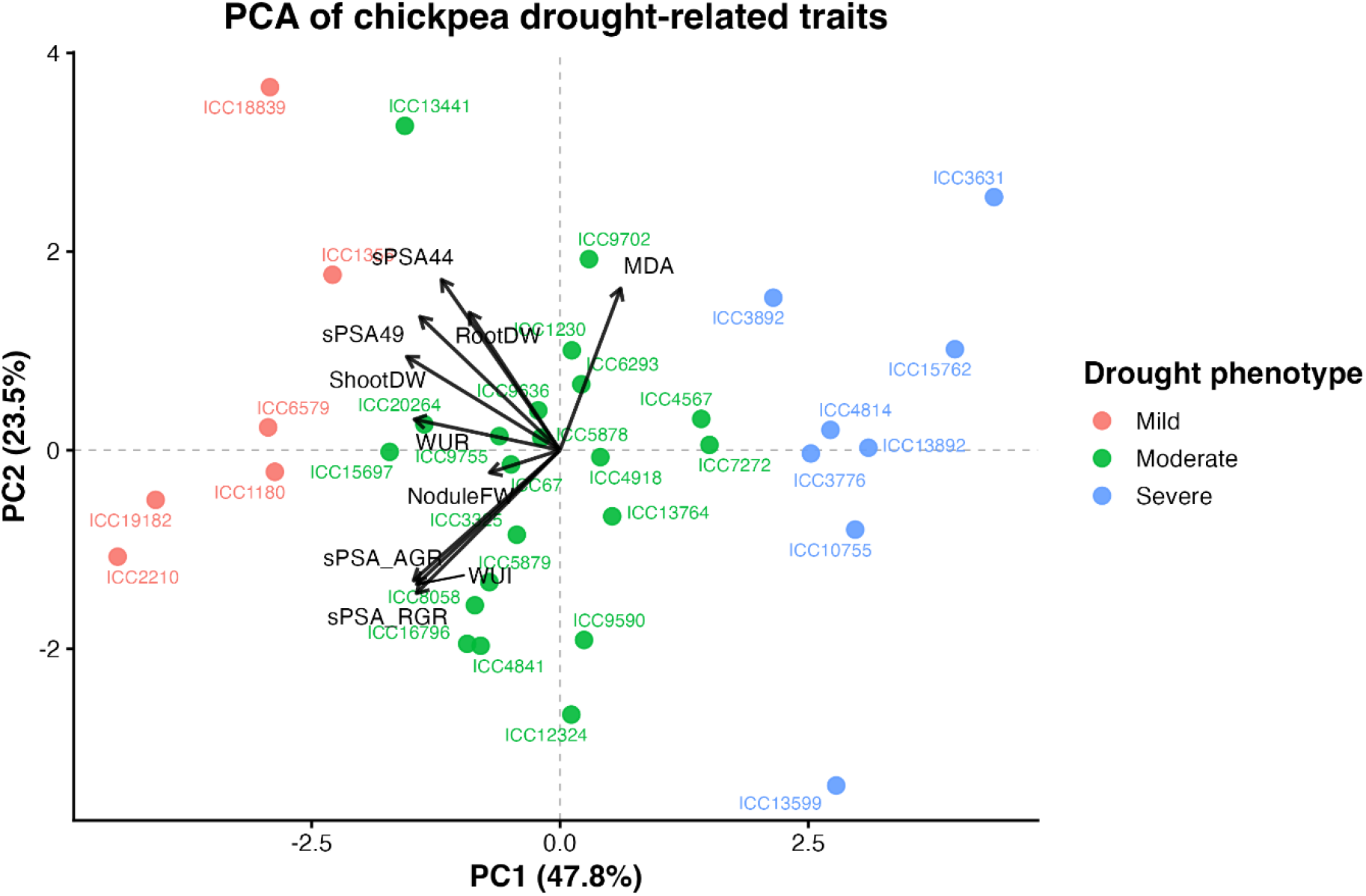
Principal component analysis (PCA) biplot of trait contributions of 35 *C. arietinum* haplotypes under two water regimes. The biplot displays the distribution of chickpea haplotypes (black points) and the contribution of the measured trait under water-limiting conditions relative to water-sufficient conditions (red vectors). Arrows indicate the direction and magnitude of trait loadings, including: nodule fresh weigh (NoduleFW); shoot dry weight (ShootDW); root dry weight (RootDW); Malondialdehyde concentration (MDA); smoothed water use rate from days 44 to 49 (WUR); smoothed water use index from days 44 to 49 (WUI); and smoothed projected shoot area (sPSA) metrics for; day 44 and 49 (sPSA44, sPSA49), average growth rate from day 44 to 49 (sPSA_AGR), and relative growth rate from day 44 to 49 (sPSA_RGR).

**Table 1:** Seed classification, malondialdehyde (MDA) and inferred tolerance of 35 *C. arietinum* haplotypes. MDA was measured from leaf tissue 14 days after drought initiation. Two leaves were harvested from each plant after imaging on 49 DAP and snap frozen in liquid N for MDA determination (n=5). Statistical significance was calculated through a two-way ANOVA with Šidák multiple comparisons test (* p≤ 0.05; ** p≤ 0.01; *** p≤ 0.001; **** p≤ 0.0001). Drought phenotype was inferred through favourable clustering of PCA1 and PCA2 in Figure 4. The ✓ symbol next to the haplotypes indicates those selected for metabolomics (Figure 5).

| HAPLOTYPE | SEED | MDA (NMOL GFW<sup>-1</sup>) |  |  |  |  | DROUGHT PHENOTYPE |
| --- | --- | --- | --- | --- | --- | --- | --- |
|  |  | Control |  | Drought |  |  |  |
|  |  | Mean | SE | Mean | SE | p |  |
| ICC67✓ | Desi | 22.0 | ± 3.31 | 36.3 | ± 5.64 | ns | Moderate |
| ICC1180 | Desi | 16.7 | ± 1.71 | 23.2 | ± 1.80 | ns | Mild |
| ICC1230 | Desi | 17.3 | ± 4.14 | 33.9 | ± 2.99 | ** | Moderate |
| ICC1356 | Desi | 18.9 | ± 1.58 | 24.6 | ± 1.89 | ns | Mild |
| ICC2210✓ | Desi | 25.2 | ± 3.32 | 27.2 | ± 2.83 | ns | Mild |
| ICC3325 | Desi | 18.3 | ± 1.12 | 28.5 | ± 1.19 | ns | Moderate |
| ICC3631✓ | Desi | 17.3 | ± 2.45 | 40.4 | ± 2.80 | **** | Severe |
| ICC3776✓ | Desi | 39.8 | ± 1.45 | 62.0 | ± 2.41 | *** | Severe |
| ICC3892 | Desi | 22.2 | ± 1.72 | 43.9 | ± 6.14 | **** | Severe |
| ICC4567 | Desi | 11.8 | ± 1.23 | 23.1 | ± 3.60 | ns | Moderate |
| ICC4814✓ | Desi | 30.9 | ± 3.46 | 53.1 | ± 3.23 | **** | Severe |
| ICC4841 | Kabuli | 18.6 | ± 2.31 | 31.0 | ± 2.93 | ns | Moderate |
| ICC5878 | Desi | 22.8 | ± 0.34 | 38.5 | ± 1.19 | * | Moderate |
| ICC5879 | Kabuli | 31.2 | ± 2.14 | 43.0 | ± 2.24 | ns | Moderate |
| ICC6293 | Desi | 38.9 | ± 4.89 | 44.7 | ± 4.19 | ns | Moderate |
| ICC6579 | Kabuli | 20.6 | ± 2.10 | 32.2 | ± 2.54 | ns | Mild |
| ICC7272 | Desi | 28.4 | ± 2.23 | 40.7 | ± 3.23 | ns | Moderate |
| ICC8058 | Desi | 33.6 | ± 2.41 | 36.3 | ± 7.23 | ns | Moderate |
| ICC9590 | Desi | 22.0 | ± 2.47 | 34.0 | ± 1.15 | ns | Moderate |
| ICC9636✓ | Desi | 28.5 | ± 2.60 | 36.1 | ± 1.97 | ns | Moderate |
| ICC9702✓ | Desi | 20.7 | ± 2.00 | 42.3 | ± 3.89 | **** | Moderate |
| ICC9755 | Desi | 25.9 | ± 1.66 | 47.4 | ± 1.92 | **** | Moderate |
| ICC10755✓ | Kabuli | 26.7 | ± 2.74 | 40.3 | ± 5.66 | ns | Severe |
| ICC12324 | Kabuli | 31.4 | ± 5.44 | 37.5 | ± 3.75 | ns | Moderate |
| ICC13441✓ | Kabuli | 18.5 | ± 3.49 | 33.6 | 2.72 | * | Moderate |
| ICC13599 | Desi | 38.0 | ± 3.00 | 38.9 | ± 2.68 | ns | Severe |
| ICC13764 | Kabuli | 25.2 | ± 1.19 | 45.0 | ± 2.29 | *** | Moderate |
| ICC13892✓ | Desi | 34.7 | ± 4.57 | 54.2 | ± 2.32 | *** | Severe |
| ICC15697 | Kabuli | 23.7 | ± 0.62 | 37.8 | ± 3.81 | ns | Moderate |
| ICC15762 | Desi | 29.7 | ± 2.06 | 45.7 | ± 7.24 | * | Severe |
| ICC16796 | Kabuli | 28.8 | ± 1.04 | 34.3 | ± 3.91 | ns | Moderate |
| ICC4918 | Desi | 24.8 | ± 1.41 | 35.2 | ± 1.44 | ns | Moderate |
| ICC19182✓ | Desi | 27.7 | ± 3.83 | 33.5 | ± 1.47 | ns | Mild |
| ICC18839✓ | Kabuli | 26.6 | ± 3.22 | 48.8 | ± 2.63 | **** | Mild |
| ICC20264 | Kabuli | 23.0 | ± 1.38 | 33.7 | ± 4.28 | ns | Moderate |

Oxidative stress was quantitated through leaf Malondialdehyde (MDA) quantification 14 days after drought initiation (49 DAP) (Table 1). MDA equivalents were greater in all water-limited plants when compared to well-watered conditions and was significant for 13 of the 35 haplotypes screened (Table 1). ICC3776, ICC4814 and ICC13892 all contained high MDA equivalents under water-stressed conditions. ICC3776, along with ICC6293 and ICC13599, also displayed high basal MDA equivalents under well-watered conditions (Table 1).

### Watering regime shifts metabolomic profile

Metabolomic profiling of leaf tissue was performed on 12 haplotypes with differing water-stress responses (Table 1, Supplementary Figure 4). PCA demonstrated a distinct separation between control and stressed treatments (Figure 5A). PC1 accounted for 39.3% of the total variance, while PC2 explained an additional 17.1%. Together, these components revealed a strong clustering pattern as metabolomic reprograming responded to well-watered or water-limited conditions.

**Figure 5:**
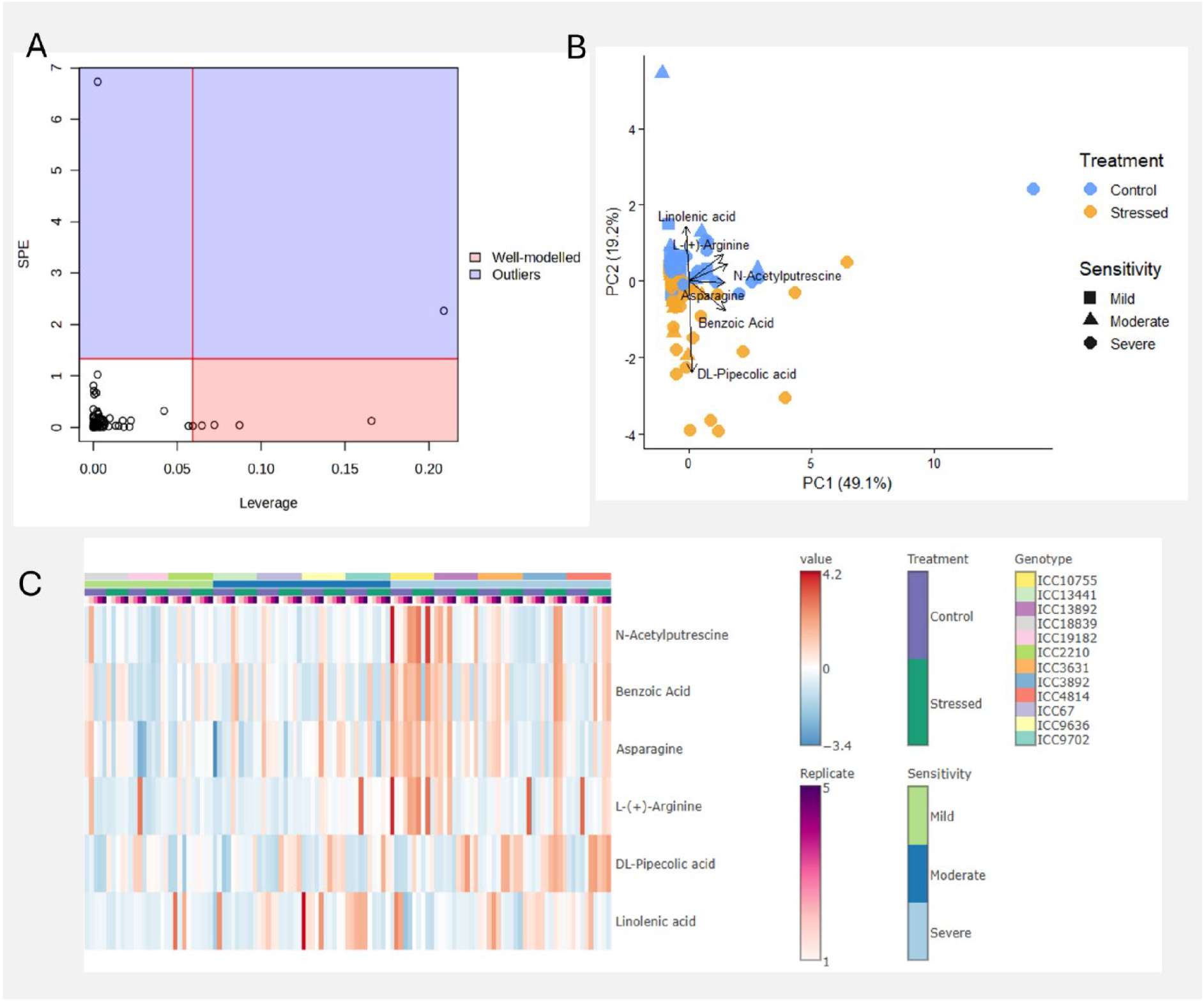
Metabolomic analysis of 12 *C. arietinum* haplotypes subjected to two water regimes. Plants were exposed to drought conditions for 14 days prior to leaf tissue sampling (n=5). (A) PCA scores plot. (B) Linear correlation analysis of treatment effect on metabolite abundance while controlling for haplotype variability, data available in Supplementary Table 1. Normalised peak heights of stress responsive metabolites (C) Succinic acid, (D) L-(-)-Proline and (E) Sucrose.

Linear correlation analysis indicated that 57 of 82 unique metabolites were significantly associated with watering regime when controlling for haplotype effects (Figure 5B). Metabolites with FDR adjusted p values below 0.05 were widely distributed across the intensity spectrum, indicating that treatment effects were not confined to only high-or low-abundance metabolites. The strong association between significance ranking and effect magnitude demonstrates a broad metabolic reconfiguration in response to stress.

Succinic acid, L-(-)-proline and sucrose displayed the most uniform response to watering regime across the 12 haplotypes investigated (Figure 5C-E). Succinic acid significantly decreased in abundance under stressed conditions, while L-(-)-proline and sucrose both increased. The observed increase in L-(-)-proline abundance was significant for all haplotypes, and this was true also for sucrose, except in ICC1075 and ICC19182.

### Metabolite drivers of the haplotype vs stress interaction

A leverage (measures a variable’s importance to the model) versus Squared Prediction error (SPE – measures goodness of fit) plot of the ASCA interaction model was generated to assess the robustness and detect influential observations (Figure 6A). Most metabolites fell within the low-leverage/low-SPE quadrant, indicating they were well modelled but uninformative. A small number of metabolites (2) exceeded threshold values for SPE and were classified as outliers. The model identified five metabolites with high leverage and low-SPE (Figure 6A). Asparagine, benzoic Acid, DL-pipecolic acid, L-(+)-arginine and N-acetylputrescine were high leverage, well-modelled metabolites associated with haplotype-specific responses to watering regime (Figure 6A). A sixth metabolite, linolenic acid, leveraged on the cutoff threshold, but when modelled along with treatment interaction displayed the highest leverage of all metabolites (leverage 0.121, SPE 2.62E-05), suggesting a strong treatment effect. The influence of these six metabolites on determining haplotype stress sensitivity was assessed through PCA (Figure 6B). Clear separation between well-watered and water-limited haplotypes was evident, and clustering of stress sensitivities was validated (Figure 6B). Haplotypes exhibiting severe stress were poorly clustered under water-limiting conditions, but those exhibiting milder stress clustered together with the well-watered controls.

**Figure 6:**
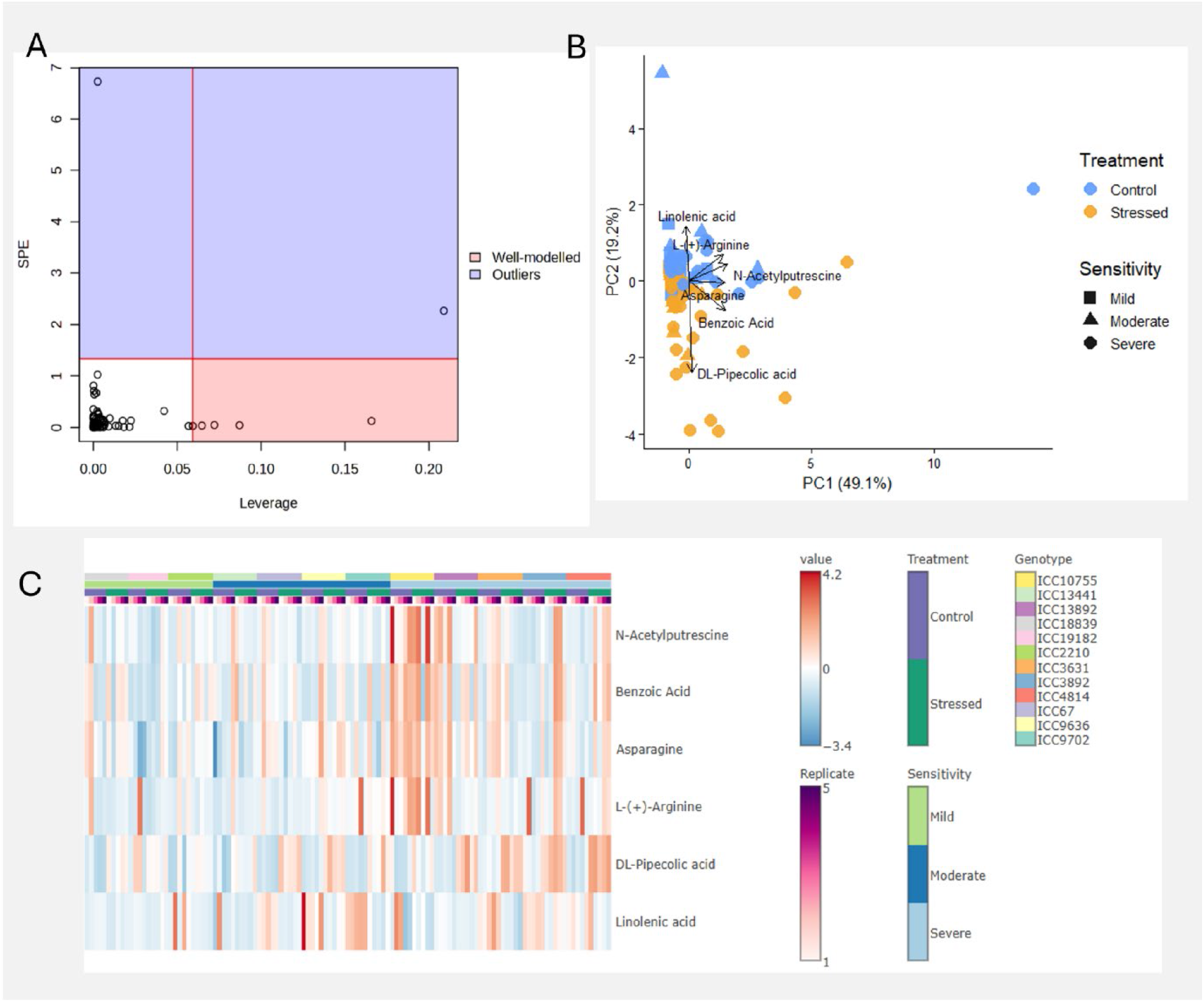
ASCA analysis highlights metabolite drivers of the haplotype × stress interaction in 12 *C. arietinum* haplotypes subjected to two water regimes. (A) Leverage vs SPE ASCA plot highlighting metabolites that are well-modelled and outliers in inferred stress sensitivity. (B) PCA of the five well-modelled metabolites driving stress tolerance. (C) Heatmap of the six well-modelled metabolites identified.

A heatmap of these six metabolites highlighted haplotype-and treatment-dependent patterns (Figure 6C). Asparagine, L-(+)-arginine, and benzoic acid had higher relative abundance under control conditions in more drought-sensitive haplotypes, with visible gradient shifts across sensitivity groups in the heatmap (Figure 6C). Meanwhile, DL-pipecolic acid accumulated in water-limited conditions, particularly with increasing drought sensitivity, while N-acetylputrescine tended to decrease in water-limited conditions, but increases were observed in stress-sensitive haplotypes. Linoleic acid abundance was more variable across haplotypes and treatments without a uniform trend.

## DISCUSSION

Chickpea is preadominantly grown under rainfed conditions, leading to periods of terminal drought during the growing season, which influences both vegetative allocation and reproductive dynamics (Hopgood et al. 2026). Drought is a problem for nitrogen-fixing species such as chickpea, where root growth is essential for the formation of rhizobial symbiosis. This study used high-throughput phenotyping and metabolomics to dissect drought responses across 35 chickpea haplotypes sourced from diversity panel of 3366 haplotypes sequenced by Varshney et al., (2021). A strong haplotype x water-regime interaction for plant water use, growth rate, nodulation and metabolomic reprogramming was observed. These findings connect haplotypes with adaptive physiological and metabolic traits that can be useful in breeding programs in the future.

### Haplotypes show diversity in water-use and growth dynamics

Plants were watered to weight and imaged on alternate days allowing clear separation of water-use trajectories between well-watered and water-limited plants after water stress imposition, with pronounced haplotypic dispersion in the shoot water use index (sWUI) across developmental windows (Figure 1). Chickpea productivity is severely impacted by drought (Anwar et al. 2025; Garg et al. 2016), contributing to a decline in chickpea yields that have stagnated at around 1 t/ha (Joshi and Rao 2017). High WUI is favourable in cropping systems, as it is a key determinant of yield (Loss et al. 1997; Siddique et al. 1998). Notably, ICC2210 and ICC18839 maintained comparatively high sWUI under both regimes, whereas ICC15762 and ICC19182 were consistently low, indicating divergent strategies for sustaining biomass under water-limiting conditions.

Genotypic variability has previously been demonstrated in chickpea: Pappula-Reddy et al. (2024b) noted that water conservation strategies of some genotypes aided post-stress physiological activities and yield. These strategies for conservative water use under drought minimise non-productive losses by selecting for high intrinsic WUE. However, Zaman-Allah et al. (2011) argue that chickpea yield under terminal drought in field conditions is not necessarily correlated with any traits related to WUE. Growth-rate estimates derived from projected shoot area declined from the onset of drought and displayed a pronounced haplotype:watering interaction. The persistence of relatively higher projected shoot area (sPSA), relative growth rate (RGR) in ICC2210 and ICC18839 under severe water limitation mirrors their favourable sWUI (Figure 2), making them ideal candidates for future breeding programs.

Nodule fresh weight was highly haplotype-dependent and drought-sensitive with nodulation of most haplotypes significantly impacted by water limitation (Figure 3). These responses are consistent with previously reported abiotic stress impacts on legume-rhizobium symbioses, where drought, salinity, heat, and nutrient imbalances can suppress infection, nodule development, and nitrogenase activity (Zahran Hamdi 1999; Lumactud et al. 2023; Fenta et al. 2012; Cerezini et al. 2017). Likewise, nitrate concentration and temperature extremes are known inhibitors of symbiotic nitrogen fixation (Ofosu-Budu et al. 1992; Streeter 1985), and for this reason were controlled for during the growing period.

Nodule weight of two haplotypes (ICC4841 and ICC15762) was not significantly impacted by watering regime, but these nodulated poorly even under well-watered conditions. ICC67 and ICC13599 both displayed favourable nodule phenotypes under water-limitation, maintaining high nodule biomass relative to their controls. Similar phenotypes have been observed in drought tolerant chickpea (Bhaskarla et al. 2020), Medicago (Staudinger et al. 2016) and soybean (Cerezini et al. 2017; Fenta et al. 2012). Our findings highlight the current diversity in nodulation and carbon allocation under water-limiting conditions in chickpea, with magnitude determined by haplotype and environment.

### Identification of favourable drought phenotypes

Mechanistic studies to define drought phenotypes of chickpea have been carried out frequently, but many of these focused on a limited number of genotypes (Garg et al. 2016; Yadava et al. 2023; Molina et al. 2008; Jain and Chattopadhyay 2010). The use of high-throughput plant phenotyping using robotic-assisted imaging platforms is necessary for large scale phenotypic screens, as manual application of water is imprecise, labour intensive and reduces imaging time (Fahlgren et a. 2015).

Recently, studies have classified genotypic tolerance to drought through both agro-physiological and biochemical characteristics (Basavaraj et al. 2025; Pappula-Reddy et al. 2024a; Pappula-Reddy et al. 2024b; Asati et al. 2024; Fazeli-Nasab et al. 2025). Furthermore, integration of ‘omic’ profiling under stress has begun to reveal the extensive reprogramming involved with abiotic stress tolerance (Duan et al. 2025; Negussu et al. 2023; Khan et al. 2019; Singh et al. 2023; Vessal et al. 2020; Kudapa et al. 2024). Here, drought sensitivity was calculated via PCA biplot of phenotypic and biochemical traits collected over the growing period (Figure 4). Clustering of favourable phenotypes classified 20% of haplotypes as mildly impacted, 54% moderately impacted and 26% severely impacted by water limitation (Table 1).

### Drought associated metabolic reprogramming

During times of low water availability, plants undergo significant acclimation, including changes in morphology, signalling, membrane stability, metabolism and osmotic adjustment (Abid et al. 2018; Basu et al. 2016; Loutfy et al. 2012). Changes in the abundance of cell metabolites is an essential process to improve water usage, protect cellular structures, maintain energy balance and prevent oxidative damage. In this study, the metabolomic shift was assessed through untargeted GC-MS, showing that typical stress responsive pathways were activated. PCA highlighted a strong treatment axis (PC1 39%) and clear separation between well-watered and water-limited treatments (Figure 5). Correlation analysis identified 57 (of 82) metabolites that were significantly correlated with watering regime across 12 contrasting haplotypes (Figure 5). Impaired energy metabolism was apparent, with some TCA intermediates prominent, including succinic acid (Figure 5). This was uniform across the panel and reveals a conserved biochemical response during times of reduced respiration and carbon supply despite haplotypic differences in growth and water use. Similar responses have previously been observed in wheat under drought stress (Bowne et al. 2012).

Reduced water availability decreases cell turgor ad disrupts normal metabolic balance, resulting in the synthesis of compatible osmolytes. Typical stress responsive osmolytes were found to accumulate, including L-(-)-proline and sucrose. Proline accumulation is a canonical drought response implicated in Osmo protection, redox poise, and stress recovery signalling (Szabados and Savouré 2010; Hanson et al. 1977). Overexpression of D-pyrroline-5-carboxylate synthetase, the rate-limiting enzyme in proline biosynthesis, does confer osmotic tolerance, at least in transgenic tobacco (Kishor et al. 1995), but there have been inconsistent reports relating proline concentrations to stress tolerance (Bandurska et al. 2017; Lv et al. 2015; Man et al. 2011; Parida et al. 2008). Consistent with these other studies, no clear correlation between drought sensitivity and proline content was observed here. Sucrose and other soluble sugars accumulated in response to water limitation (Figure 5), but this is typical under water limitation, where osmoregulation is key in cell protection (Chaturvedi et al. 2024; Sehgal et al. 2018; Du et al. 2020; Morgan 1984). It is also possible that accumulation of sucrose in leaves is a result of impaired phloem transport causing a build-up in photosynthetic tissues. More recently, the role of sucrose synthase 1 has been elucidated in *Arabidopsis* mutants under drought, with T-DNA mutants showing a reduction in soluble sugar levels and increase in MDA, thereby highlighting the contribution to drought-induced osmotic adjustment (Gurrieri et al. 2020).

### Metabolomic profiling for drought-tolerance

ASCA and follow up modelling highlighted six metabolites that correlated strongly with haplotype: watering interaction, including Asparagine, Benzoic Acid, DL-Pipecolic acid, L-(+)-Arginine, Linoleic acid and N-acetylputrescine (Figure 6). Asparagine has been shown to accumulate in response to diverse stressors and is thought to have an essential role in nitrogen remobilisation (Oddy et al. 2020). Yadav et al. (2019) suggested that asparagine accumulation negatively correlates with yield, and this aligns with the findings of the present study, where asparagine relative abundance under control conditions increased with drought sensitivity (Figure 6). Similarly, L-(+)-arginine is involved in nitrogen transport and storage, as a precursor for the synthesis of polyamines and nitric oxide (Shi et al. 2013; Tun et al. 2006). In this study, under control conditions, L-(+)-arginine pools were notably lower across stress tolerant haplotypes, possibly due to higher nitric oxide and polyamine pools which could prime cells to tolerate water-limitations. In *Arabidopsis*, arginase mutants lower arginine pools by directing metabolism to polyamines and nitrous oxide, and in-turn increase tolerance to multiple abiotic stressors (Shi et al. 2013). Benzoic acid followed a similar trend to asparagine and L-(+)-arginine, but in general was more stress responsive across the sensitivity index. Benzoic acids are important building blocks for primary and secondary metabolites, including key hormones such as salicylic acid which confers stress tolerance to multiple abiotic stressors (Senaratna et al. 2003; Widhalm and Dudareva 2015). Exogenous application of benzoic acid has also been reported to improve gas exchange and chlorophyll retention under drought in soybean (Anjum et al. 2013).

Accumulation of DL-Pipecolic acid was strongly associated with tolerance, increasing with drought sensitivity (Figure 6). Interestingly, pipecolic acid is more strongly associated with systemic acquired resistance under biotic rather than abiotic stressors (Kumari et al. 2025; Koc and Seckin Dinler 2022). However, its accumulation in response to osmotic potential in rapeseed (Moulin et al. 2006), and protective roles in drought through interaction with antioxidant systems in tomato (Wang et al. 2021), have been reported. DL-pipecolic acid provides a novel metabolomic marker for osmotic stress in chickpea. N-acetylputrescine is another novel marker identified in this study; in stress tolerant haplotypes it was highly responsive to watering regime. This may reflect a mechanism to maintain polyamine homeostasis and avoid stress induced damage during water-limitation (González-Hernández et al. 2022), but requires further investigation to fully elucidate these pathways. Functionally, these metabolite patterns correlate with traits - haplotypes maintaining higher sWUI and sPSA RGR under drought (e.g., ICC2210, ICC18839, ICC19182) tended to express metabolite changes consistent with osmotic adjustment and signalling mediated acclimation, whereas haplotypes with large growth penalties showed stronger oxidative signatures that could be further elucidated by integrating time resolved metabolomics with dynamic water use metrics.

## CONCLUSION

This study highlights substantial haplotypic variation in chickpea responses to drought, integrating physiological parameters with metabolomic traits. Clear divergence in water-use dynamics and growth under stress revealed strong haplotype:watering interactions, with ICC2210, ICC18839 and ICC19182 haplotypes maintaining favourable water-use efficiency and growth, identifying them as promising drought-tolerant candidates. Drought reduced nodulation in most haplotypes, but some retained symbiotic performance, indicating possible exploitable resilience. Metabolomic profiling showed widespread reprogramming under stress, including reduced TCA intermediates and increased osmoprotectants such as proline and sucrose. Key metabolites (Asparagine, Benzoic Acid, DL-Pipecolic acid, L-(+)-Arginine, Linoleic acid and N-acetylputrescine) were linked to haplotype-specific stress responses, reflecting differences in nitrogen metabolism and stress signalling. These metabolites provide a foundation for targeted breeding for improved chickpea productivity under terminal drought.

## Data availability

The data that support the findings of this study are available from the corresponding author upon reasonable request.

## Acknowledgements

We thank the staff at The Plant Accelerator (Waite campus, Adelaide University) a National Collaborative Research Infrastructure Strategy (NCRIS) facility for phenotyping of chickpea haplotypes.

## Author contributions

S.A.R., C.S and D.D. helped with funding acquisition. S.A.R. and N.B. designed experiments, supervised the research, analysed data along with A.C. N.B. and S.A.R. drafted the manuscript. All authors had intellectual input into the project and commented on the manuscript.

