## Supplementary figures for "Integrated phenotypic and metabolomic analyses identify elite chickpea haplotypes under water limitation": Supplementary Figures.pdf

| Lane \ Position | 4 | 5 | 6 | 7 | 8 | 9 | 10 | 11 | 12 | 13 | 14 | 15 | 16 | 17 | 18 | 19 | 20 | 21 | 22 | 23 |
|---|---|---|---|---|---|---|---|---|---|---|---|---|---|---|---|---|---|---|---|---|
| 2 | L18 Str | L18 Ctrl | L30 Ctrl | L30 Str | L16 Ctrl | L16 Str | L01 Str | L01 Ctrl | L22 Str | L22 Ctrl | L11 Str | L11 Ctrl | L33 Ctrl | L33 Str | L04 Ctrl | L04 Str | L14 Ctrl | L14 Str | L30 Str | L30 Ctrl |
| 3 | L33 Ctrl | L33 Str | L35 Ctrl | L35 Str | L07 Ctrl | L07 Str | L17 Str | L17 Ctrl | L29 Ctrl | L29 Str | L20 Ctrl | L20 Str | L06 Ctrl | L06 Str | L29 Str | L29 Ctrl | L35 Ctrl | L35 Str | L23 Ctrl | L23 Str |
| 4 | L25 Str | L25 Ctrl | L05 Ctrl | L05 Str | L26 Ctrl | L26 Str | L23 Str | L23 Ctrl | L11 Ctrl | L11 Str | L09 Ctrl | L09 Str | L10 Ctrl | L10 Str | L13 Ctrl | L13 Str | L24 Str | L24 Ctrl | L34 Str | L34 Ctrl |
| 5 | L19 Str | L19 Ctrl | L02 Str | L02 Ctrl | L20 Str | L20 Ctrl | L10 Ctrl | L10 Str | L24 Ctrl | L24 Str | L18 Ctrl | L18 Str | L28 Str | L28 Ctrl | L26 Ctrl | L26 Str | L32 Ctrl | L32 Str | L27 Str | L27 Ctrl |
| 6 | L13 Ctrl | L13 Str | L12 Ctrl | L12 Str | L27 Ctrl | L27 Str | L14 Ctrl | L14 Str | L08 Ctrl | L08 Str | L03 Str | L03 Ctrl | L05 Ctrl | L05 Str | L16 Str | L16 Ctrl | L31 Str | L31 Ctrl | L17 Ctrl | L17 Str |
| 7 | L06 Ctrl | L06 Str | L32 Ctrl | L32 Str | L34 Ctrl | L34 Str | L04 Str | L04 Ctrl | L31 Ctrl | L31 Str | L07 Str | L07 Ctrl | L08 Ctrl | L08 Str | L21 Ctrl | L21 Str | L22 Str | L22 Ctrl | L02 Ctrl | L02 Str |
| 8 | L21 Ctrl | L21 Str | L09 Ctrl | L09 Str | L15 Ctrl | L15 Str | L03 Ctrl | L03 Str | L28 Str | L28 Ctrl | L12 Str | L12 Ctrl | L01 Ctrl | L01 Str | L15 Str | L15 Ctrl | L25 Str | L25 Ctrl | L19 Str | L19 Ctrl |
| 9 | L28 Str | L28 Ctrl | L12 Ctrl | L12 Str | L33 Str | L33 Ctrl | L22 Str | L22 Ctrl | L34 Ctrl | L34 Str | L08 Ctrl | L08 Str | L18 Str | L18 Ctrl | L26 Str | L26 Ctrl | L04 Ctrl | L04 Str | L35 Ctrl | L35 Str |
| 10 | L35 Ctrl | L35 Str | L01 Str | L01 Ctrl | L23 Str | L23 Ctrl | L31 Ctrl | L31 Str | L21 Ctrl | L21 Str | L32 Str | L32 Ctrl | L02 Str | L02 Ctrl | L09 Str | L09 Ctrl | L29 Str | L29 Ctrl | L01 Str | L01 Ctrl |
| 11 | L04 Str | L04 Ctrl | L24 Ctrl | L24 Str | L27 Str | L27 Ctrl | L15 Str | L15 Ctrl | L05 Ctrl | L05 Str | L25 Ctrl | L25 Str | L16 Str | L16 Ctrl | L14 Str | L14 Ctrl | L28 Str | L28 Ctrl | L20 Ctrl | L20 Str |
| 12 | L19 Str | L19 Ctrl | L08 Ctrl | L08 Str | L03 Str | L03 Ctrl | L32 Str | L32 Ctrl | L16 Str | L16 Ctrl | L10 Str | L10 Ctrl | L33 Ctrl | L33 Str | L07 Str | L07 Ctrl | L21 Str | L21 Ctrl | L27 Ctrl | L27 Str |
| 13 | L11 Ctrl | L11 Str | L06 Ctrl | L06 Str | L13 Str | L13 Ctrl | L07 Str | L07 Ctrl | L02 Ctrl | L02 Str | L34 Ctrl | L34 Str | L11 Ctrl | L11 Str | L05 Ctrl | L05 Str | L17 Str | L17 Ctrl | L22 Ctrl | L22 Str |
| 14 | L30 Str | L30 Ctrl | L26 Str | L26 Ctrl | L29 Str | L29 Ctrl | L10 Str | L10 Ctrl | L14 Ctrl | L14 Str | L23 Ctrl | L23 Str | L31 Ctrl | L31 Str | L19 Ctrl | L19 Str | L12 Str | L12 Ctrl | L13 Str | L13 Ctrl |
| 15 | L09 Str | L09 Ctrl | L17 Str | L17 Ctrl | L25 Str | L25 Ctrl | L20 Str | L20 Ctrl | L18 Ctrl | L18 Str | L30 Ctrl | L30 Str | L15 Str | L15 Ctrl | L24 Ctrl | L24 Str | L06 Ctrl | L06 Str | L03 Str | L03 Ctrl |
| 16 | L08 Ctrl | L08 Str | L15 Ctrl | L15 Str | L34 Str | L34 Ctrl | L01 Ctrl | L01 Str | L29 Str | L29 Ctrl | | | | | | | | | | |
| 17 | L20 Str | L20 Ctrl | L13 Ctrl | L13 Str | L30 Str | L30 Ctrl | L21 Ctrl | L21 Str | L05 Ctrl | L05 Str | | | | | | | | | | |
| 18 | L16 Ctrl | L16 Str | L35 Str | L35 Ctrl | L12 Ctrl | L12 Str | L11 Str | L11 Ctrl | L10 Str | L10 Ctrl | | | | | | | | | | |
| 19 | L23 Str | L23 Ctrl | L28 Str | L28 Ctrl | L07 Ctrl | L07 Str | L24 Str | L24 Ctrl | L18 Str | L18 Ctrl | | | | | | | | | | |
| 20 | L09 Str | L09 Ctrl | L14 Ctrl | L14 Str | L31 Str | L31 Ctrl | L27 Ctrl | L27 Str | L06 Ctrl | L06 Str | | | | | | | | | | |
| 21 | L04 Str | L04 Ctrl | L02 Ctrl | L02 Str | L03 Str | L03 Ctrl | L25 Ctrl | L25 Str | L22 Ctrl | L22 Str | | | | | | | | | | |
| 22 | L33 Ctrl | L33 Str | L17 Str | L17 Ctrl | L26 Str | L26 Ctrl | L32 Ctrl | L32 Str | L19 Ctrl | L19 Str | | | | | | | | | | |

**Supplementary Figure 1:** The randomized layout of the experiment on the conveyor system in the NE Smarthouse. A split-unit design was used to randomly allocate the 35 haplotypes (L01-L35) to main units and the two Waterings (Ctrl = Control; Str = Stressed) to the two carts within a main unit. The blue lines encompass the five blocks, each of which contains one replicate of the 70 combinations of haplotypes and waterings; the purple lines demarcate the main units and the plot cells are coloured for the two carts within a main unit.

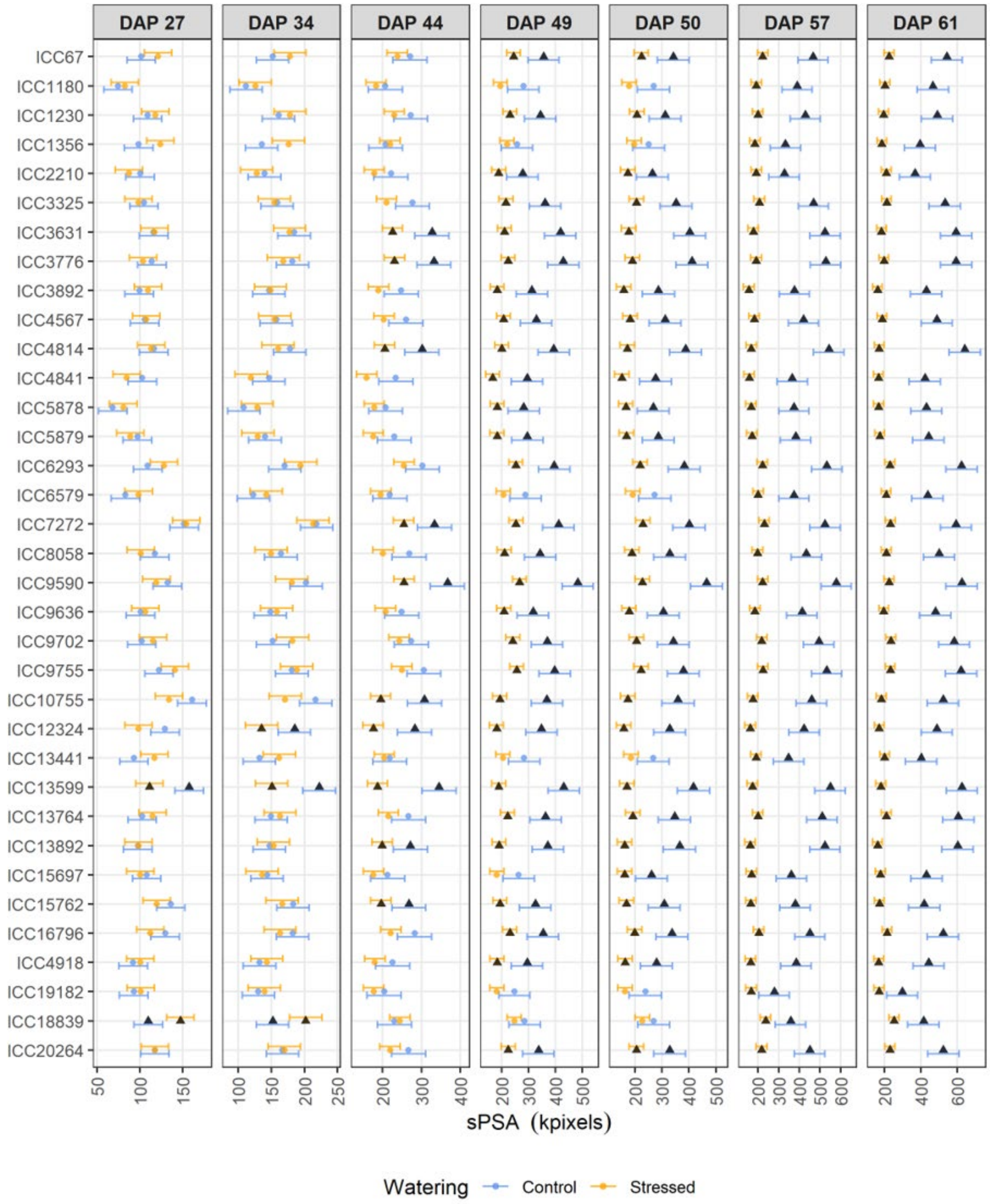


**Supplementary Figure 2:** The smoothed Projected Shoot Area (sPSA) for 35 haplotypes subjected to two water regimes. Error bars are an EMM (n = 5) $\pm$ half-LSD (5%). The EMMs from the same DAP and Watering level are significantly different ($p \leq 0.05$) if their error bars do not overlap. Haplotypes for which the difference between the watering treatments is significant ($p \leq 0.05$) are plotted with a solid triangle for both treatments.

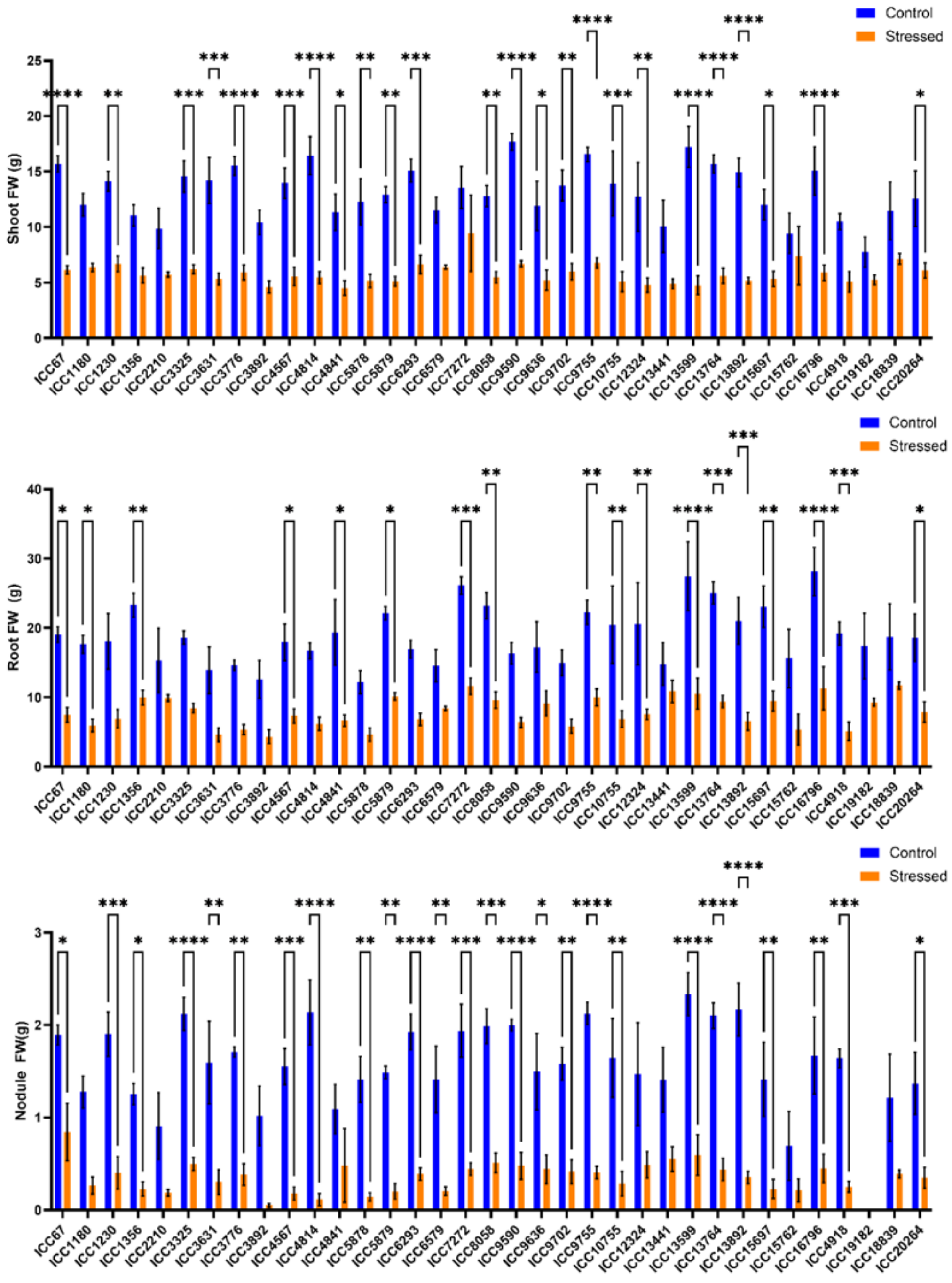


**Supplementary Figure 3:** Effect of drought stress on shoot, root, and nodule fresh weight of chickpea haplotypes. (A) shoot fresh weight (FW), (B) root fresh weight (FW), and (C) nodule fresh weight (FW) of 32 chickpea haplotypes grown under control and stressed conditions and harvested 63 days after planting. Data are mean ± standard error (SE). Asterisks indicate significant differences between control and drought treatments for each genotype ($P < 0.05$, $P < 0.01$, $P < 0.001$, $P < 0.0001$) based on two-way ANOVA with Šídák's multiple comparisons test.

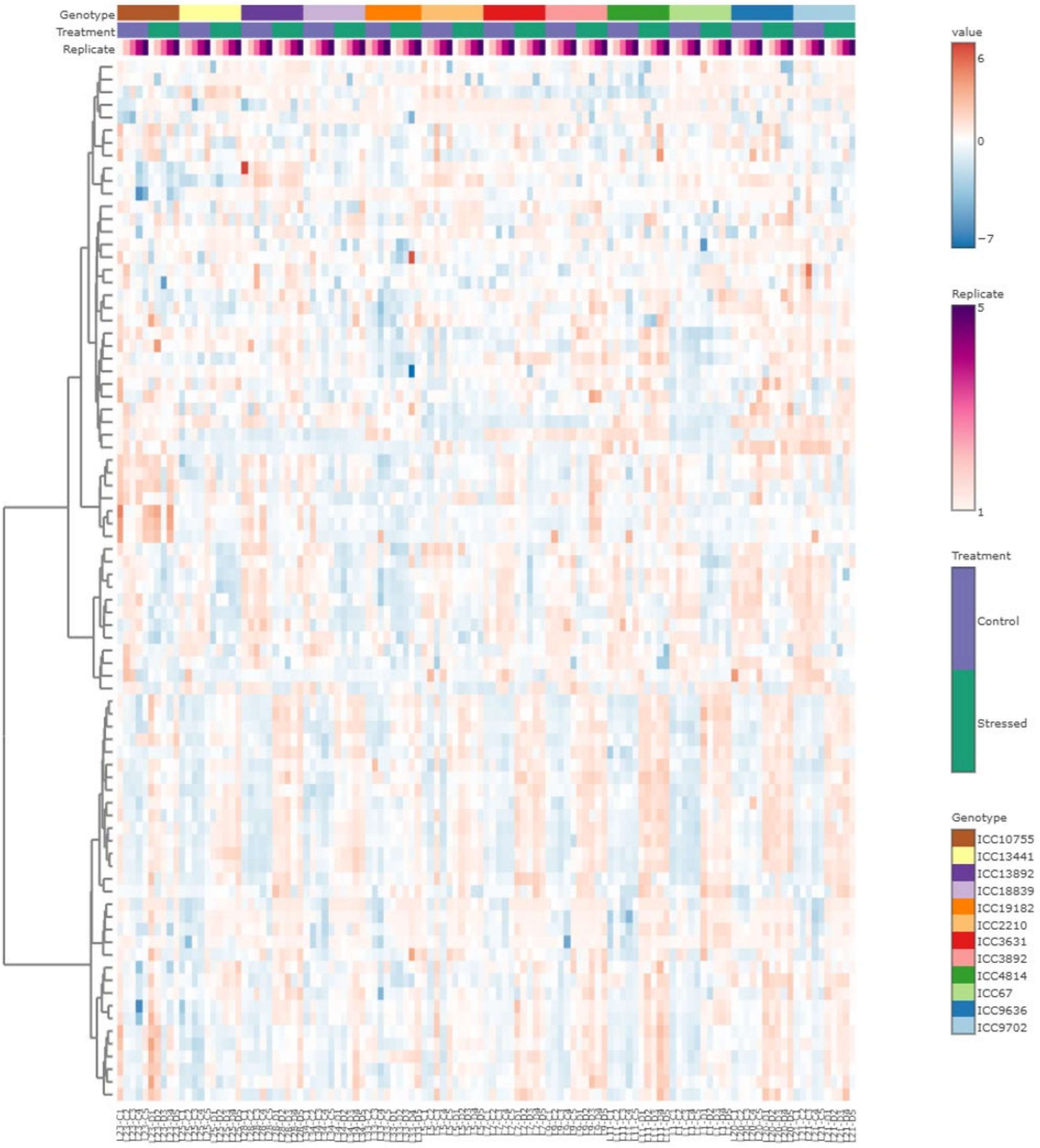


**Supplementary Figure 4:** Heatmap of the 82 identified metabolites identified in 12 *C. arietinum* haplotypes subjected to two water regimes. Plants were exposed to drought conditions for 14 days prior to leaf tissue sampling (n=5).
